# Weakly-bound Dimers that Underlie the Crystal Nucleation Precursors in Lysozyme Solutions

**DOI:** 10.1101/275222

**Authors:** M.C. Byington, M.S. Safari, V. Lubchenko, J.W. McCabe, L.A. Angel, D.H. Hawke, S.J. Bark, J.C. Conrad, P.G. Vekilov

## Abstract

Protein crystallization is central to understanding of molecular structure in biology, a vital part of processes in the pharmaceutical industry, and a crucial component of numerous disease pathologies. Crystallization starts with nucleation and how nucleation proceeds determines the crystallization rate and essential properties of the resulting crystal population. Recent results with several proteins indicate that crystals nucleate within preformed mesoscopic protein-rich clusters. The origin of the mesoscopic clusters is poorly understood. In the case of lysozyme, a common model of protein biophysics, earlier findings suggest that clusters exist owing to the dynamics of formation and decay of weakly-bound transient dimers. Here we present evidence of a weakly bound lysozyme dimer in solutions of this protein. We employ two electrospray mass spectrometry techniques, a combined ion mobility separation mass spectrometry and a high-resolution implementation. To enhance the weak but statistically-significant dimer signal we develop a method based on the residuals between the maxima of the isotope peaks in Fourier space and their Gaussian envelope. We demonstrate that these procedures sensitively detect the presence of a non-covalently bound dimer and distinguish its signal from other polypeptides, noise, and sampling artefacts. These findings contribute essential elements of the crystal nucleation mechanism of lysozyme and other proteins and suggest pathways to control nucleation and crystallization by enhancing or suppressing weak oligomerization.

## INTRODUCTION

Protein crystallization is an integral component of processes in nature, industry, and the laboratory. Proteins are stored as compact crystals in viruses, bacteria, and seeds (1, 2). On the other hand, formation of crystals and other ordered solids of wild-type or mutant proteins contributes to the pathology of condensation diseases, such as age-onset nuclear cataracts (3, 4), Alzheimer’s and other neurological conditions (5, 6), and sickle cell anemia (7, 8). Crystalline preparations offer advantages as vehicles for sustained drug release. With the significant number of proteins now applied as medication, protein crystallization is a vital part of the pharmaceutical industry (9-11). Lastly, understanding of protein function relies on accurate representations of the three-dimensional arrangement of the atoms in a protein molecule and the majority of protein structures today are determined by x-ray crystallography (12-14). Progress in understanding the natural processes and controlling the pathological and engineered ones in these and other areas relies on improved understanding of protein crystallization.

Crystallization starts with nucleation, in which the formation of new crystal embryos in a supersaturated solution is inhibited by the free energy loss due to the creation of a crystal-solution interface (15-17). Nucleation represents the rate-limiting stage of crystallization and determines the main properties of the crystal population, including the crystal polymorph, the number of crystals, and their size distribution. A protein solution supersaturated with respect to a crystal overcomes the nucleation barrier by means of localized fluctuations that bring both the solute concentration and detailed particle arrangement close to those in the incipient crystal (17). According to classical nucleation theory, the rarity of successful transitions over the barrier strongly delays nucleation and constrains the overall crystallization rate (18). Surprisingly, recent experimental measurements of nucleation rates revealed that they are even *lower*, by many orders of magnitude, than those predicted by theory (19-21). The issue of low nucleation rates and several other unexplained features of protein nucleation kinetics was resolved by the discovery that the nuclei of ordered protein solids form not in the supersaturated solution but rather in preexisting mesoscopic protein-rich clusters (22-30). In addition to crystals, the mesoscopic clusters host the nucleation of other ordered protein solids, such as sickle-cell hemoglobin polymers (27) and amyloid fibrils (28-30).

Mesoscopic clusters have been demonstrated in solutions of numerous proteins at various pHs, ionicities, temperatures, and compositions (26 31-35). Their diameters are of order 100 nm (22, 26, 34, 35) and each cluster contains 10^4^ – 10^5^ protein molecules (26, 32-34, 36, 37). With lysozyme, a robust protein used as a model in several biophysics fields, cluster formation is reversible and the clusters exchange protein with the host solution (33 37-40). Remarkably, the responses of the characteristic cluster size and phase volume to variations of the ionic strength, pH, and additive concentration are *decoupled* and the cluster size is independent of protein concentration, solution ionic strength, and pH (39, 40). These findings are in sharp contrast to observations for typical phase transitions, in which the size of the incipient domains and the volume occupied by the emerging phase increase or decrease concurrently (41). They suggest that the formation mechanism of the mesoscopic clusters is fundamentally different from those underlying other protein condensates.

**SCHEME 1.**
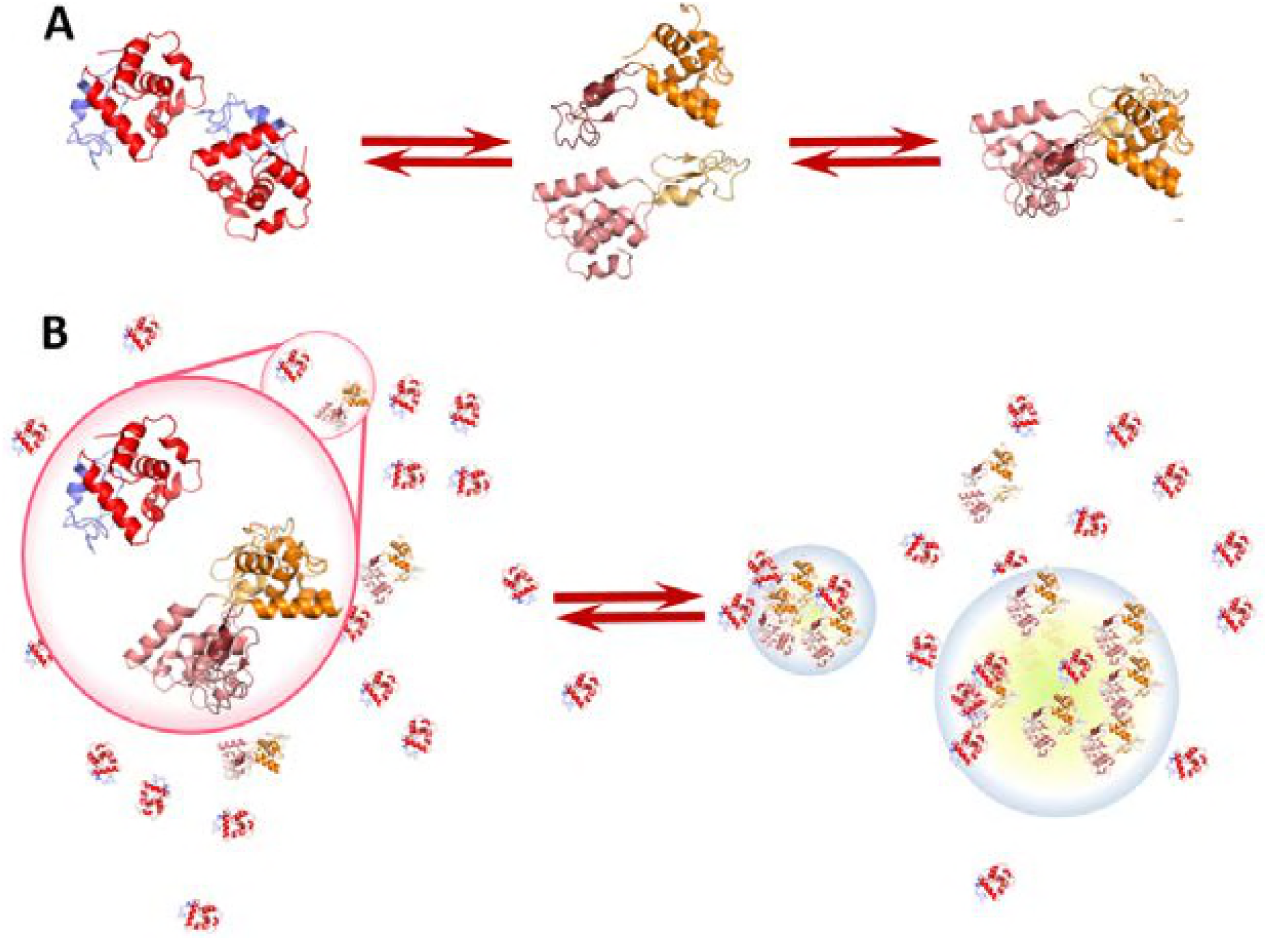
The proposed mechanism of the mesoscopic lysozyme clusters. **(A)** The formation of transient dimers bound by hydrophobic contacts between the peptide backbones exposed after partial unfolding. **(B)** The segregation of dimers and monomers into clusters.

Exploring the puzzling cluster behaviors revealed that whereas the volume occupied by the clusters population is dictated by the thermodynamics of the protein solution, the size of the clusters reflects the dynamics of formation and decay of protein complexes (32, 37, 39, 40). The complexes must be weakly-bound and decay after a certain lifetime (32, 33, 40, 42). They may be dimers, trimers, etc. for single-chain protein, or misassembled oligomers of multiple-chain proteins (23, 43). Characterization of the cluster populations in lysozyme solutions after destabilization of the protein conformations with urea (40, 44) and in sheared solutions (39, 45) suggest that enhanced partial unfolding of lysozyme molecules is a part of the mechanism of lysozyme clusters. Such unfolding exposes hydrophobic surfaces between the lysozyme constituent domains to the aqueous solution and facilitates the formation of domain-swapped dimers, Scheme 1a. NMR tests of conformational instability as a part of the lysozyme cluster mechanism revealed that the flexibility of lysozyme molecules held in clusters in much greater than of the molecules residing in a dilute solution (44, 46), consistent with partial unfolding along the hinge between the two lysozyme structural domains, identified in (47). These results support the notion that partial unfolding of the two lysozyme domains is a part of the mechanism of formation of lysozyme mesoscopic clusters, illustrated in Scheme 1. To date, however, dimers have not been directly detected.

Here we report the detection of a lysozyme dimer, whose existence underlies the mechanism of formation of the mesoscopic lysozyme-rich clusters illustrated in Scheme 1, by electrospray ionization mass spectroscopy (ESI-MS) (48). Because of the ability of electrospray to volatilize large biomolecules (49), ESI-MS has become a vital tool in the study of protein biophysics (50). Of the currently available ionization methods, ESI-MS is viewed as the least destructive and least likely to produce artefacts not present in solution (51-56). The technique is particularly advantageous in investigations of non-covalent protein interactions (52) since the employed solvent evaporation times are significantly shorter than the diffusion times of molecular species and even fast diffusion-limited reactions fail to proceed (56). ESI-MS has produced valuable data on metal binding to proteins (57), dimerization and oligomerization (58), and the conformation of protein complexes (57, 59, 60), among others. To detect and identify a lysozyme dimer present at low concentration, we design and implement a new method to analyze high-resolution mass-spectra in the Fourier domain.

## MATERIALS AND METHODS

### Solution preparation

Lysozyme from hen egg white (Thermo Scientific #89833) used for these experiments was dissolved in DI water (18.2 MΩ by Millipore MilliQ Gradient) containing 20 mM ammonium acetate and pH was adjusted to 7.8. The solution was dialyzed against 1 L of the solvent using dialysis cassettes with 2000 g mol^-1^ cutoff (Life Technologies Corp #66203) for 38-48 hours in a refrigerator (4°C) on a stir plate at ∼50 rpm with a 5 cm magnetic stir bar. After dialysis, the solution was filtered through syringe filters with 220 nm pore size (LightLabs TC-9747). The concentration was determined by diluting an aliquot of the prepared solution 1000-fold and measuring the light absorption at 280 nm using a Beckman DU-800 spectrophotometer.

### Electrospray ionization mass spectroscopy (ESI-MS)

In electrospray ionization, solution droplets are ejected from a small nozzle (with diameter from less than a micron to 100 micron) by applying a high voltage between the nozzle and the entrance to a chamber filled with inert gas at low pressure. The droplets flying into this chamber shrink as the solvent evaporates. The shrinking concentrates the dissolved ions, which increases the Coulomb energy of a droplet until it fractures. Evaporation, charge compression, and explosion repeat until the solvent is fully evaporated and the dissolved ions are suspended in the gas phase (53, 56, 61). Electrospray ionization has several advantages to other methods of sample dispersion applied in mass spectroscopy: it is less likely to fragment a molecule or a molecular complex; it does not require extensive sample preparation or large sample volumes; ions are generated directly from the solution of interest, and, most importantly, it allows for observation of molecules in their solution states (55, 56, 62). ESI has been demonstrated to transfer proteins from the solution to the gas while conserving their mass, charge state, binding interactions, and conformation (63, 64).

We employed two methods of ESI-MS. We used a combined ion mobility separation mass spectrometry (IMS-MS) Waters Synapt high-definition G1 device equipped with an ESI source and a quadrupole–ion mobility–orthogonal time-of-flight configuration (65). IMS-MS simultaneously measures the mass-to-charge ratios (*m*/*z*) and arrival times that relate to the identity, interactions, and conformation of the protein ion. High resolution mass spectra were obtained using an Orbitrap Fusion Tribrid Mass Spectrometer from ThermoFisher Scientific with a parallel electrospray nozzle. This instrument combines quadrupole, ion trap and Orbitrap mass analysis in a Tribrid architecture to achieve ultrahigh resolution (66).

### Immunoblotting identification of the protein species in lysozyme solutions

We prepared a 4-20% gradient gel by polymerizing bisacrylamide and acrylamide in the presence of tetramethylethylenediamine and ammonium persulfate; the four reagent were purchased from BioRad. We used a 2× Laemmli solution (4% sodium dodecyl sulfate, 10% 2-mercaptoethanol, 20% glycerol, and 0.125 M Tris-HCL at pH 6.8), purchased from Sigma-Aldrich, as a running buffer. Transfer to a polyvinylidene fluoride (PVDF, BioRad) membrane was carried out for 16 hours at 16 V potential in a transfer buffer composed of 20% (v/v) methanol, 25 mM Tris, 190 mM Glycine, and 0.05% SDS in water. Detection was achieved using a hen egg white lysozyme antibody HyHEL-5, generously provided by the R.C. Willson group (67), horse radish peroxidase, and 3,3’,5,5’-tetramethylbenzidine (TMB), both from Sigma-Aldrich.

## RESULTS AND DISCUSSION

### Characterization of lysozyme solutions by ion mobility separation mass spectroscopy

Tests of the presence of weakly bound transient dimers of lysozyme were carried out in solutions with pH 7.8, equal to that in numerous studies of the mesoscopic lysozyme-rich clusters (32, 33, 38-40, 44, 45, 68). We use the ammonium acetate buffer; it is preferable for the present study to the often employed 4-(2-hydroxyethyl)-1-piperazineethanesulfonic (HEPES) buffer, which may cause ion suppression in the electrospray ionization chamber (69, 70). Ion mobility separation mass spectroscopy (IMS-MS) produces a two-dimensional spectrum, where the signal from each ion is correlated with its drift time and the mass to charge ratio, *m/z*, Fig. 1A. Integrating over the drift time yields a standard mass spectrum correlation between signal intensity and the *m/z*ratio, Fig. 1B.

**FIGURE 1.**
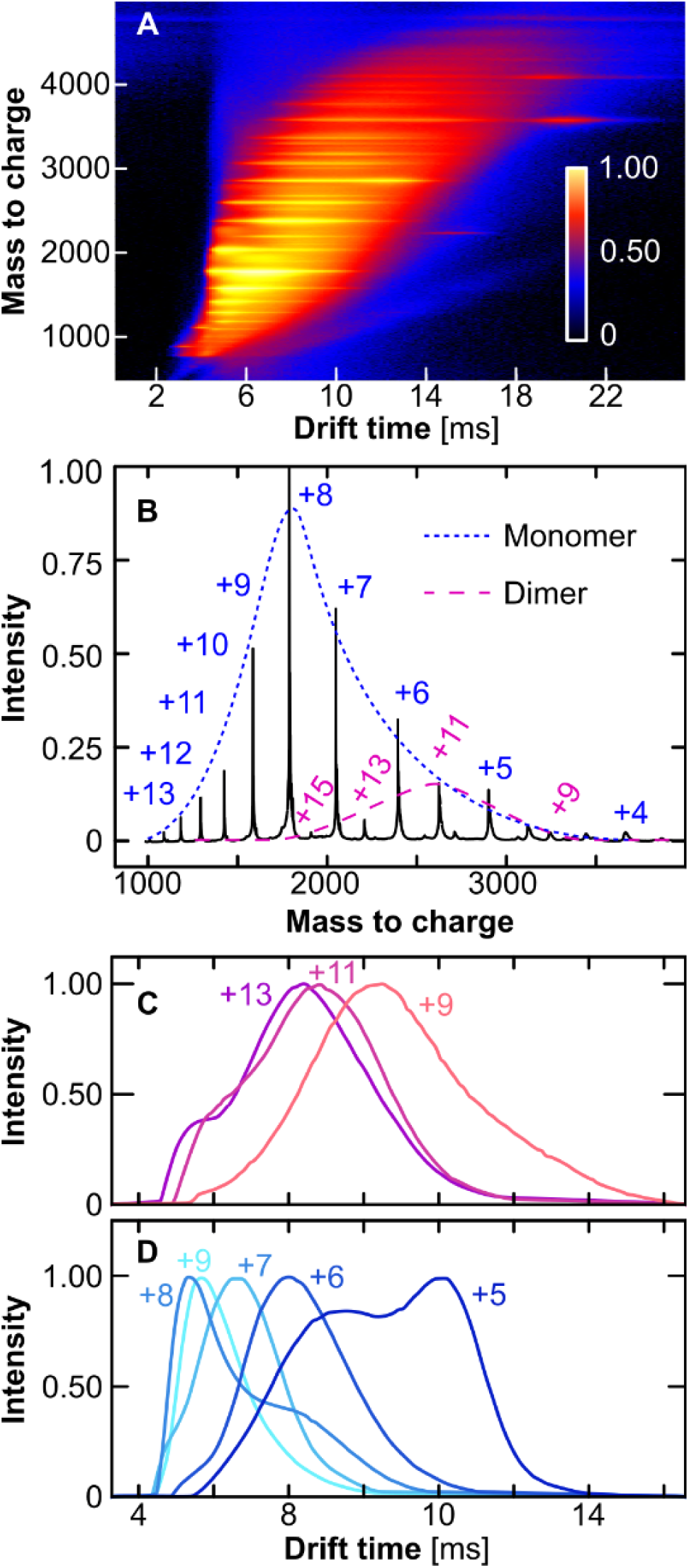
Ion mobility mass spectra of lysozyme. **(A)** Intensity as a function of mass to charge ratio and drift time. **(B)** Mass spectrum integrated over drift time. The monomer and dimer charge state distributions, shown as dashed lines, reveal that the dimer distribution is shifted to lower charges, corresponding to higher *m/z*ratios. **(C)** and **(D)** The drift time distributions of dimers and monomers, respectively, at several *m\*values. Solution pH was 7.8 and concentration was 1.1 mg/mL = 80 μM in all panels.

The molecular weight of lysozyme of 14.3 kDa and the range of output *m/z* ratios, from 1000 to 5000, constrain the charges of the detected monomers from +13 to +4. Protein molecules attain charge owing to protonation of basic aminoacid side chains, arginine, lysine, and histidine, or deprotonation of the acidic residues glutamate and aspartate. At the tested pH 7.8, the balance of 17 positive and nine negative groups equals an average net charge of eight (42, 71). Lysozyme monomers with +8 charges generate the strongest signal in Fig. 1, A and B. The correspondence between the most populated state in the mass spectrum and the molecular charge in the solution suggests that the electrospray dispersion does not modify the average charge of the molecules.

The spectra in Fig. 1 A and B are consistent with the presence of weakly bound dimers in the tested solution. A dimer bound by non-covalent bonds has exactly twice the mass of a monomer (covalent binding may involve cleavage of water, H_2_, etc., or the association of bridging residues, with the corresponding loss or gain of mass) and twice the number of residues that may be charged due to protonation or deprotonation. Hence, in the *I(m/z)* plots in Fig. 1B the peaks of a dimer in even charge states would exactly overlap the monomer peaks with half that charge, making the monomer and dimer signals indistinguishable. Because fractional charge states are impossible, dimers with odd charge states will be unobstructed by monomers. The mass spectra in Fig. 1, A and B reveal a population of lysozyme dimers with odd charges between +15 and +9. Comparing the area under the Gaussian envelope of the dimer peaks to that under envelope of the monomer peaks suggests that the dimers account for about 10% of the intensity of the signal between *m/z* = 1000 and 4000. The intensity of the dimer signal is not correlated to the fraction of dimers contained in the sample since different species may elicit dramatically distinct responses in the mass spectroscopy signal. The signal of the dimer is shifted to higher *m/z* ratios, corresponding to lower charge *Z*. This shift indicates that the dimer may be weakly bound and decay owing to repulsion between highly charged constituent monomers.

The drift times of the odd-charged dimers in Fig. 1C and monomers and even-charged dimers in Figs. 1D increase with decreasing charge. Similar increases in the literature have been attributed to charge effects on both the apparent cross sectional collision area and the electromotive force in the drift tube (58 72-74). The distribution of the drift times of monomers and dimers with even-charges in Figs. 1D and S1 reveal an apparently monomodal distribution for monomers with charge +9 and bimodal distributions corresponding to monomers with charges +8, +7, +6, and +5. The drift times of the slower components of the four bimodal distributions are longer than those of the faster component by ca. 1.45 – 1.6, suggesting that they represent dimers with double molecular weight and charges +16, +14, +12, and +10; such dimers would diffuse with 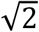 slower characteristic velocities because of their greater mass. The absence of a slow mode in the peak corresponding to the +9-charge monomer indicates inhibited formation of dimers with charge +18, likely owing to Coulomb repulsion between similarly charged monomers. The distributions of the drift times of the odd-charged dimers in Fig. 1D are broader and in some case, bimodal. These distributions are consistent with two or more dimers present in the chamber whose distinct mobilities may be associated with different collision cross-sections (75).

In further support of the notion that dimers with charges in the range from +16 to +10, suggested by the ion mobility distributions and mass spectra in Fig. 1, represent weakly-bound complexes, the signals from dimers with charges +9 and +15 disappear when the concentration of the tested solution is lowered to 5 μM, Fig. S1. By contrast, the signal from monomers with charge +4, equal in intensity to that of dimers with +9 and +15 charges at 80 μM, is still strong. Solvent evaporation in the electrospray chamber may induce charge states not present in the original tested solution (56). The characteristic times of evaporation, ion condensation, and droplet explosion are, however, shorter than the characteristic times of diffusion (56) and are likely to prevent artefacts due to the synthesis of dimers during ion dispersion.

### High-resolution mass spectra

Further tests of the presence of weakly-bound dimers in lysozyme solutions employed a high-resolution mass spectrometer from ThermoFisher. This instrument outputs mass spectra in the *m/z* range from 1400 to 1800 Da, Fig. 2A, which contain signals from lysozyme monomers with charges +10, +9, and +8. To analyze whether the spectra contain signals from dimers with *z* in the range from +20 to +16, we developed a model for the isotope composition of the signal, readily resolved by this instrument. The premise underlying the data analysis is that lysozyme represents a mixture of molecules, whose molecular weight varies by integer multiples of 1 Da owing to the isotopic variety of the constituent elements. Thus, a high-resolution monomer mass spectrum would consist of peaks corresponding to individual isotopes separated by 1/*z* and clustered in three groups separated by charge as displayed in Fig. 2 B, D, and F. The mass of a dimer also changes in discrete steps of size 1 Da and the peaks from the individual dimer isotope configurations would be separated by 1/*z*. The charge of the dimers, however, is double that of the monomers, and the dimer peaks are periodically distributed at half the *m/z* step of the monomer. Thus, the presence of dimers in the solution would have two manifestations in the mass spectrum: emergence of extra peaks between the monomer isotope peaks and enhanced intensity of the monomer peaks. To identify the weak modifications of the spectrum due to the dimers, we compare experimentally recorded mass spectra of lysozyme to a model spectrum of pure monomers. To amplify the spatially-periodic dimer signal, we work in the Fourier domain.

**FIGURE 2.**
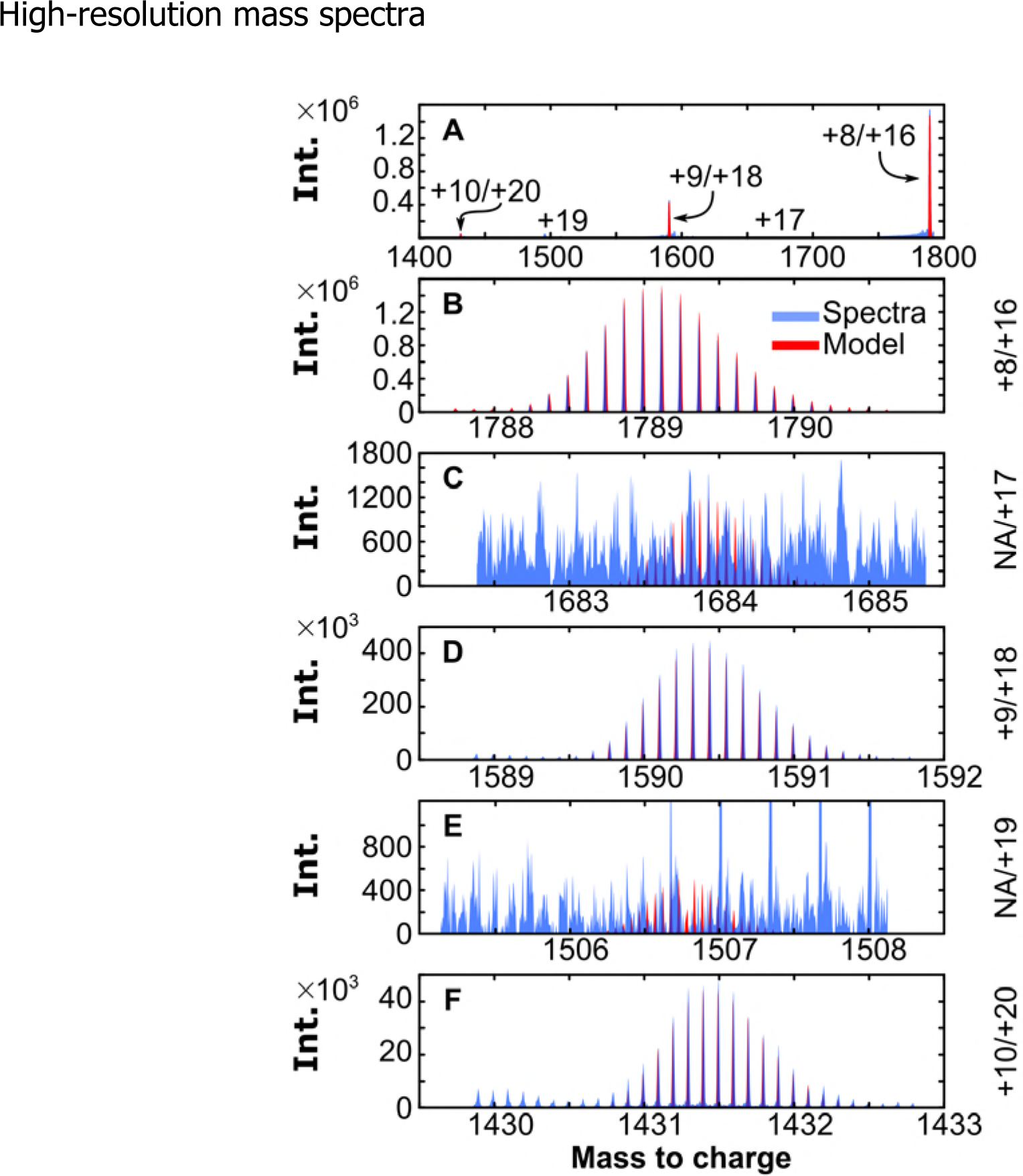
High resolution mass spectra of 40 mg/mL solution. **(A)** The entire spectrum. The locations of signals from monomers/dimers with their respective charges indicated. **(B)** – **(F)**. Zoom-ins of five *m/z* regions. The charges of the monomers/dimers that could produce signal in the respective *m/z* ranges are displayed at right. Models of monomer spectra accounting for the isotope composition are superimposed in red in B, D, and F; models of dimer spectra, in C and E.

### Fourier analyses of high-resolution mass spectra of monomer-dimer mixtures

To model the isotope distribution of lysozyme, we follow the polynomial method (76, 77). We represent the chemical formula of lysozyme, H_959_C_613_S_10_O_185_N_193_, as 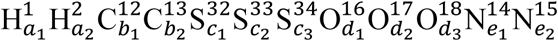. The latter expression accounts for the two primary isotopes of hydrogen, H^1^ and H^2^, and two, three, three, and two, respectively, of carbon, sulfur, oxygen, and nitrogen; the superscripts are equal to the mass number of each isotope; the subscripts designate the number of atoms of each isotope and sum to the respective total number of atoms of each element, e.g., for H, *a*_*1*_ *+ a*_*2*_ *= 959*. For each element, the probability of a particular isotope distribution is given by a relation similar to that for H

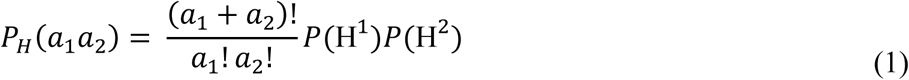

where *P(*H^*n*^) is the abundance of the hydrogen isotope with a mass number *n* (78). To evaluate the probability of a specific isotopic configuration of lysozyme, we use that the isotope distribution of an element is independent of the others and obtain

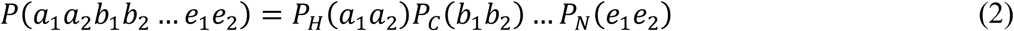

Equation (2) predicts billions of isotope combinations. The majority of these, however, are rare and represent a minor fraction of the lysozyme molecules. For instance, the rarest isotope configuration would occur naturally in one out of 10^3000^ molecules; the latter number is greater than the estimated number of atoms in the known universe, about 10^82^. We neglect the unlikely combinations and evaluate the probabilities of about 10^6^ isotope configurations. We obtain a discrete probability distribution *P(m/z)*, where *m* is the sum of the isotope masses (different than the mass number (78)) of a given isotope configuration and *z* is the molecular charge.

To account for the experimental uncertainty and the unequal contributions of added neutrons to the masses of different elements (79), we replace each *P(m/z)* datum point with a Gaussian curve of width σ, centered at the respective *m/z* value. We obtain a continuous expression for the probability of a *m/z* value

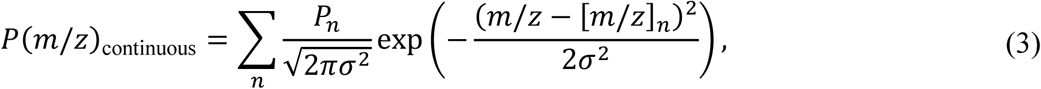

where *n* distinguishes the isotope configurations, [*m/z*]_*n*_ is the discrete *m/z* value of the *n*-th configuration, and *P*_*n*_ is its probability resulting from Eq. (2). The selection of *σ* is informed by the resolution of the experimental data.

Assuming that the mass spectrum signal at a *m/z* value is proportional to the its probability

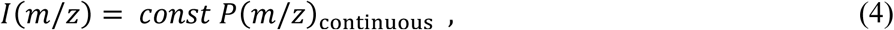

this model accurately reproduces the experimental mass spectra of lysozyme in the +8, +9 and +10 charge states, Fig. 2 B, D, and F. The centers of the model peaks diverge from those in the mass spectrum by less than 0.005 Da, Fig. S2, commensurate with the calibration accuracy of the mass spectrometer. Comparison of the model prediction for the isotope distribution of dimers with charges +17 and +19 with the mass spectrum reveals no detectable signal from dimers with these charges, Fig. 2 C and E.

In the Fourier domain, groups of *I(m/z)* peaks with equal *z*, defined in Eqs. (3) and (4), transforms into *I(ω)* (ω is the Fourier frequency with dimension *z/m*), which consist of a similar sequence of peaks, see Supporting Material for details. Analyses of *I(ω)* reveal that it contains a set of narrow peaks modulated by a relatively broad envelope. The envelope over *I(ω)* peaks that belong to one charge group represents the Fourier transform of the shape of the individual *I(m/z)* peaks. Since the Fourier transform of a Gaussian function is another Gaussian function, the envelope over *I(ω)* peaks is Gaussian regardless of the relative magnitudes of the *I(m/z)* peaks. The width of an individual *I(ω)* peak is the inverse of the width of the *I(m/z)* envelope, whereas the width of the envelope of the *I(ω)* peaks is the inverse of the width of an individual peak in *m/z* space, up to a factor of order one. Owing to the double charge of the dimers, the spacing Δω between the dimer peaks in *I(ω)* is twice as broad as that between monomer peaks; this ratio is reciprocal to the ratio between the spacing 1/*z* between the monomer and dimer peaks in *I(m/z)*, which is smaller by half for the dimers. Most importantly, since the signal from monomers and dimers is recorded with the same instrument calibration constant, the spectra of the monomers and dimers are additive. We use these four properties of *I(ω)* to probe whether high-resolution mass spectra of lysozyme reveal the presence of weakly-bound dimers.

We evaluated the impact of the presence of dimers on *I(ω)* and its distinction from the inevitable noise in experimentally measured *I(m/z)*. Using the polynomial method, we computed the mass spectrum of a +8 charged lysozyme monomer, Fig 3A. To test the manifestations of uncontrolled noise on the Fourier transform of the mass spectrum, we superimposed random fluctuations, Fig. 3B. A third model spectrum was composed by adding the spectrum of a dimer with signal 99× weaker than that of the monomer, Fig. 3C. We computed the Fourier transforms of all three spectra, Fig. 3 D, E, and F. We fitted the maxima of the peaks in the three resulting Fourier distributions with Gaussian curves. Analyses of the mass spectra in the Supporting Material suggest that that the envelope of the maxima of monomer Fourier peaks should represent a Gaussian function and this correlation is independent of the relative magnitudes of the *I(m/z)* peaks. Subtracting the maximum values of the Fourier peaks from the respective values of the best Gaussian fit generates the residuals displayed in Fig. 3G – I. In agreement with these analyses, these residuals are close to zero for the pure monomer model, Fig. 3G. The small deviations near the two edges of the ω field are attributable to the limited *m/z* range of the spectrum in Fig. 3A.

**FIGURE 3.**
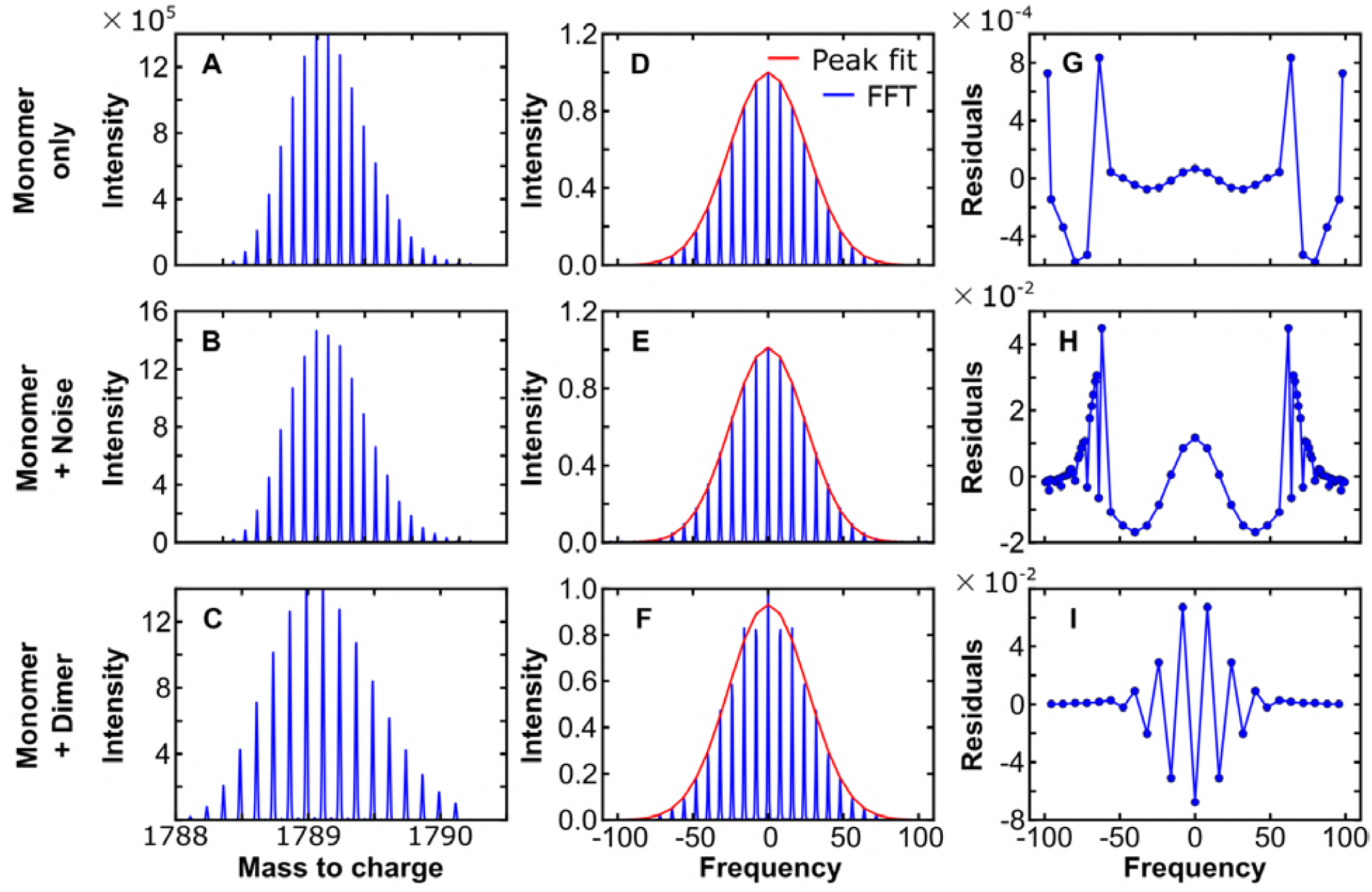
Analyses of model mass spectra of monomer/dimer mixtures in the Fourier domain. **(A)** Simulated mass spectrum of lysozyme monomer with +8 charges. **(B)** Same as in a with added random noise. **(C)** Same as in a with an added dimer signal weaker by 99×. **(D)** – **(F)** Fast Fourier transforms of respective spectra in a – c. The peak maxima were fitted with a Gaussian function, displayed in red. **(G)** – **(I)**. The residuals of the Fourier amplitudes computed by subtracting the best-fit Gaussian from the values of the peak maxima in, respectively, D – F.

Adding noise to the monomer signal increases the magnitude of the residuals by two orders of magnitude, particularly at high-frequency values (Fig. 3H). The residuals from the combined monomer and dimer signals display a characteristic saw-tooth pattern Fig. 3I), which manifests the addition of dimer intensity to every other monomer Fourier peak. As demonstrated in the analyses in the Supporting Material, the separation in ω space between the dimer peaks is twice that between the monomer peaks. The amplification of the periodic dimer signal in the Fourier space enables the detection of dilute dimer in the presence of noise. The characteristic saw-tooth pattern in the residuals between the Fourier amplitudes and their Gaussian fit values is detectable if the intensity of the dimer signal is greater than the fluctuations caused by the finite sampling interval, illustrated in Fig. 3G.

### Detection of a weakly-bound lysozyme dimer

We applied the insights on the signature patterns of the Fourier peaks residuals over their Gaussian envelopes arising from the above model to the +8 and +9 charge segments of the high-resolution lysozyme mass spectra. The data obtained from a solution with concentration 40 mg/mL produce a saw-tooth pattern that nearly coincides with the pattern resulting from a model mass spectrum assuming that the dimer produces 1 % of the total signal intensity, Fig. 4 A – C. The correspondence between the measured and the model spectra is consistent with the presence of dimers with charge +16 in this solution. We carried out identical analyses of the +9 charge segment of the mass spectrum at the same concentration. The plot of the Fourier peak residuals, Fig. 4D, does not exhibit the saw-tooth pattern and is akin to the pattern resulting from the superposition of random noise on the monomer mass spectrum, suggesting that dimers with charge +18 are not present in the solution. This observation is consistent with the mass spectra in Figs 1 and 2, which indicate the presence of dimers of charge up to +16 and the absence of dimers of charges +17 and +18.

**FIGURE 4.**
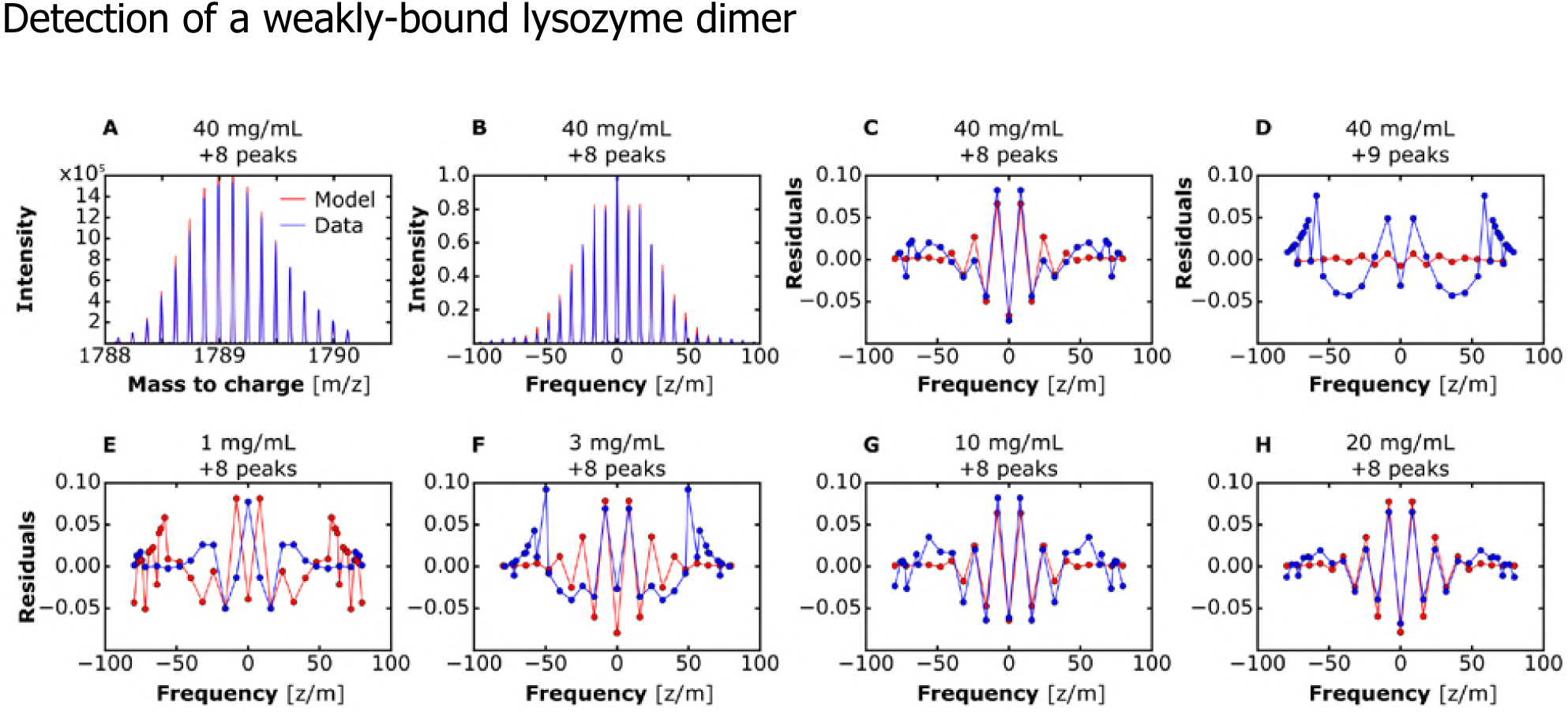
Detection of lysozyme dimers in experimental mass spectra. **(A)** Mass spectrum of lysozyme solution at 40 mg/mL in the *m/z* region of +8 charges. A simulated mass spectrum of a mixture of 99/1 monomer/dimer is superimposed in red. **(B)** Fourier transforms of measured and simulated spectra. **(C)** – **(H)** The residuals of a Gaussian function fitted to the maxima of the Fourier peaks over the values at these maxima. **(C)** For the solution concentration and charge state characterized in A and B. **(D)** – **(H)** For solution concentrations and charge states as indicated on each panel.

To monitor the correlation between the strength of the dimer signal and the total protein concentration, we measured the mass spectra of solutions with concentrations 1, 3, 10 and 20 mg/mL. We isolated the +8 charge segment for each concentration, and computed its Fourier transform and the corresponding Fourier peak residuals, Fig. 4 E – H. For each concentration, we compared the pattern of residuals with that of a model assuming the dimer produces 1 % of the total signal intensity, Fig. 4 E – H. For the 1 mg/mL solution, the residuals of the measured spectrum are distinct from those of the model, suggesting the absence of dimers at this concentration. The saw-tooth pattern of the residuals plot for the 3 mg/mL solution is consistent with a weak dimer signal, whereas the patterns for solutions with 10 and 20 mg/mL exhibit the dimer signature. The absence of signal from +18 charged dimers and the weak dimer signal at low total protein concentrations suggest that the dimer indicated by the +8 charge segment of the monomer mass spectrum for solutions with concentration above 10 mg/mL is weakly bound and decays if highly charged or diluted. These conclusions are consistent with the inferences arising from the EMS-MS data in Fig. 1.

### Covalent or weakly-bound dimers

To test if the dimer that induces the modifications in the monomer mass spectra, discussed above, is covalently-bound, we modeled the modification of the mass spectra peaks due to such dimers. A covalent dimer that results from a condensation reaction, e.g., 2Lys → Lys^2^ + H_2_O, or is assembled by bridging, would attain a different molecular mass. Thus, the release of a water molecule with molecular weight 18 Da would shift the group of peaks with +16 charges to *m/z* values lower by 1.125 Da. The peaks from the +8 charged monomers and +16 charged dimers extend over about 2 Da, hence, the contribution of a condensation dimer to this group of peaks would be significantly shifted to lower *m/z* values, Fig. 5A. By contrast, the contribution of a weakly-bound dimer with +16 charges and molecular weight exactly double that of the monomer would be uniformly distributed over the peaks of the monomer with charge +8, Fig. 5B.

**FIGURE 5.**
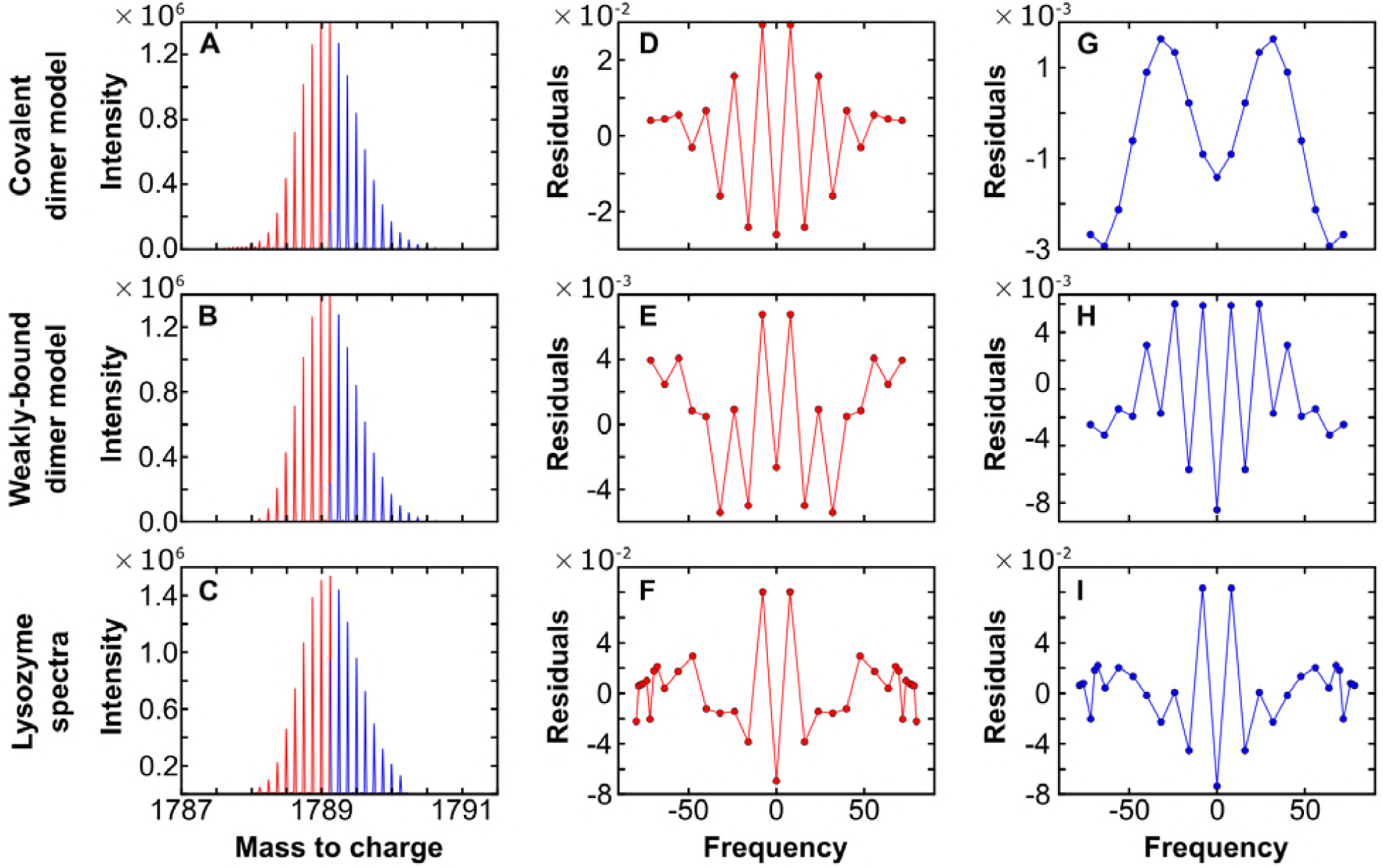
The manifestation of covalently-bound dimers. **(A)** and **(B)** Modeled spectra of +8 charged monomer in the presence of a dimer with +16 charges. We assume the dimer molecular weight is lower by 18 Da from the double molecular weight of the monomer in A and is exactly double in B. **(C)** The +8 charges segment of the mass spectrum of lysozyme at 40 mg/mL. **(D)** – **(I)** Residuals of a Gaussian function fitted to the maxima of the Fourier peaks over the values at these maxima. In **(D)** – **(F)**, we considered the Fourier transforms of the lower molecular weight peaks highlighted in red in, respectively, A – C; in **(G)** – **(I)**, of the peaks with higher-molecular weight, highlighted in blue.

To establish whether the dimer contribution to the spectrum of the +8 charged monomers is symmetric or asymmetric, we analyzed independently the high and low molecular weight segments of the spectrum, highlighted in blue and red, respectively, in Fig. 5 A – C. We modeled how the presence of covalent and weakly bound dimers would modify the patterns of the residuals between the best Gaussian fit and the Fourier peak maxima of either the left or the right half of this mass spectrum, Fig. 5 D, G, E, and H. The modeling results reveal that a dimer with lower molecular weight would generate a saw-tooth pattern from the low-molecular weight segment of the mass spectrum (Fig. 5D), but will only produce a pattern similar to finite sampling range artefacts (Fig. 3G) on the high molecular weight side. By contrast, a weakly-bound dimer with molecular weight exactly double that of the monomer generates modified saw-tooth patterns with both the left and right halves of the spectrum (Fig. 5 E and H). Analogous analysis of the measured spectrum of a lysozyme solution (Fig. 5C) reveals saw-tooth patterns resulting from both low and high molecular weight mass spectrum data, Fig. 5 F and I. These tests indicate that the dimer detected here has molecular weight within 6 Da (determined by the resolution of the mass spectrum) of the double mass of a lysozyme monomer and is likely bound by weak non-covalent interactions.

### Immunoblotting tests for the presence of other proteins

Previous analyses of commercial lysozyme preparations have suggested the presence of up to five additional proteins with molecular weights 18, 28, 39, 66, and 78 kDa (80-82). Whereas the 28 kDa heterogeneity was tentatively identified as a lysozyme dimer produced by oxidation, the identity of the other polypeptides was not firmly established (80). This uncertainty leaves open the possibility that the specie contributing to the mass spectra in the *m/z* range from 1787 to 1791 Da might be, e.g., a protein with molecular weight 39,358 Da, which carries +22 charges. To test this hypothesis, we examined the purity of the employed lysozyme preparation by sodium dodecyl sulfate polyacrylamide gel electrophoresis (SDS PAGE). We tested the identity of the found extraneous proteins by immunoblotting and their mechanism of formation by applying an oxidation potential, Fig. 6.

**FIGURE 6.**
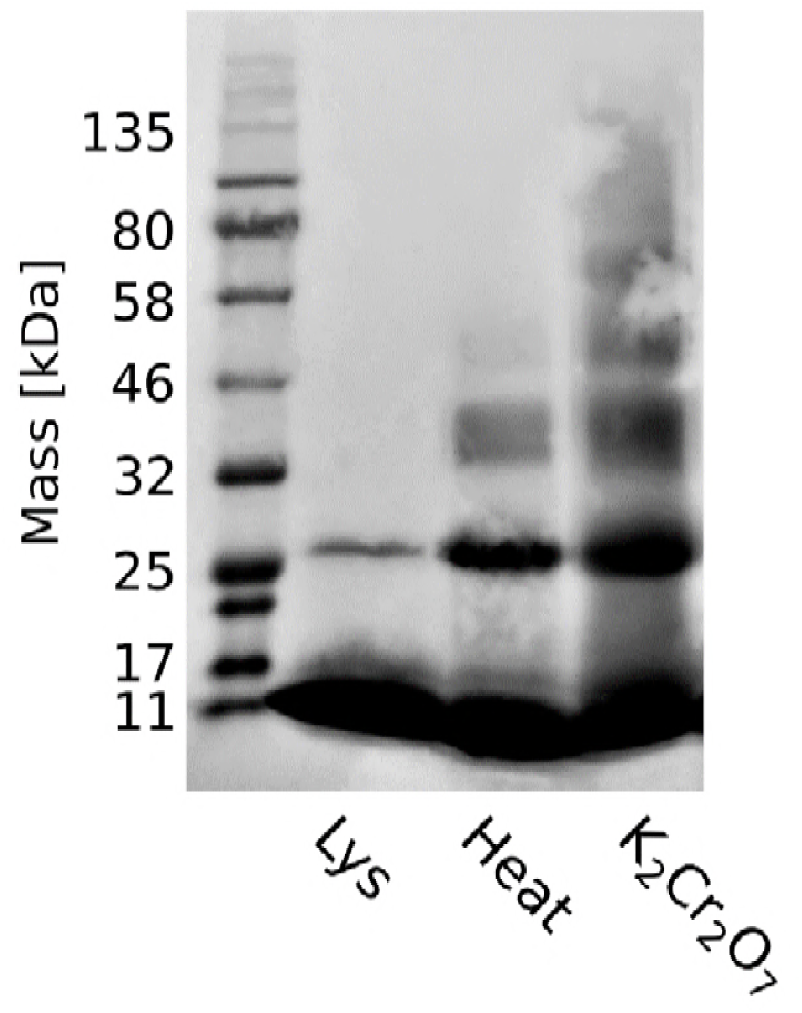
Immunoblot of a sodium dodecyl sulfate polyacryl amide electrophoresis gel (SDS-PAGE) to identify the protein species in lysozyme solutions. The four gel lanes display a pre-stained molecular weight ladder, a lysozyme solution at 5 mg/mL, the same solution after exposure to 100°C for 15 minutes, and in the presence of 1 mM K_2_Cr_2_O_7_.

The second lane of the gel displayed in Fig. 6 reveals that the employed lysozyme preparation contains a single heterogeneity. This protein has a molecular weight of ca. 27 kDa, roughly double the molecular weight of native lysozyme, and produces a strong signal when treated with an antibody for hen egg white lysozyme. In accord with previous identifications (80-82), the immunoblotting response implies that this is a lysozyme dimer. Sodium dodecyl sulfate fully denatures all treated proteins, hence, a weak bond between the constituents of this lysozyme dimer is unlikely. To establish the mechanism of its formation, we subjected the solution to oxidative stress in two ways: we exposed solution aliquots either to overheating at 100°C for 15 minutes (high temperature accelerates oxidation by soluble O_2_) or to 1 mM K_2_Cr_2_O_7_, a strong oxidizing agent. Both treatments produced several other species with approximate molecular weights 42, 55, and 70 kDa. The strong binding of the lysozyme antibody to these bands indicates that they are three, four, and five member oligomers of lysozyme, formed in a redox reaction. Importantly, the applied oxidative stress strongly enhances the intensity of the dimer band. This observation suggests that the 27 kDa lysozyme dimer and the other observed oligomers are produced in redox reactions. Such polypeptides likely have distinct molecular weights and charge states than expected for an exact weakly-bound dimer and their signals in the mass spectrum are likely out of the ranges probed in Figs. 1 and 2.

## CONCLUSIONS

We characterized the protein composition of a solution of highly purified lysozyme by two electrospray mass spectrometry techniques: a high-resolution implementation and a combination with ion mobility spectrometry. To enhance the high-resolution signal from potential secondary species in the high-resolution mass spectra, we developed a method based on the residuals between the maxima of the isotope peaks in Fourier space and their Gaussian envelope. Both methods reveal signals consistent with a lysozyme dimer with molecular weight exactly double that of the monomer, present at low concentration. Analyses of the sensitivity of the residuals method indicates that any covalently bound dimers, whose molecular weight deviates from that of two monomers by more than 6 Da, would generate a distinct residuals pattern. The dimer signal vanishes at high charges of the detected lysozyme ions and at low protein concentrations, suggesting the decay of a weakly-bound charged complex. Immunoblotting analyses of the tested protein solutions indicates that the known covalently bound lysozyme oligomers are produced by redox reactions and their molecular weight and charges place them out of the ranges of the analyzed mass spectra. These findings demonstrate the presence of weakly-bound transient dimers in lysozyme solutions, which underlie the existence of mesoscopic protein-rich clusters (32, 33, 39, 40, 44).

Even though experimentally measured mass spectra are adequately modeled by assuming that 1% of the signal is generated by dimers, the varying sensitivity of the detectors to different analyzed species prohibits any inferences on the fraction of dimers in the tested lysozyme samples; it is likely that the dimer fraction is significantly lower than 1%. Furthermore, the data presented here provide no evidence regarding the structure of the detected weakly-bound dimers or their spatial distribution throughout the solution volume. A series of published results, however, demonstrates that native lysozyme molecules do not dimerize (42) and partial unfolding that exposes to the solution the hydrophobic interface between the constituent structural domains drives the unfolded molecules into mesoscopic protein-rich clusters (38-40, 44). Collectively, the published and present results suggest that the dimers consist of partially unfolded lysozyme molecules bound by hydrophobic contacts between their interdomain interfaces, as illustrated in Scheme 1A, and assemble into mesoscopic clusters, Scheme 1B (32).

The mesoscopic clusters serve as preferred sites for the nucleation of crystals composed of native monomers (22, 23, 44), indicating that despite the accumulation of dimers, the clusters are rich in monomers. The co-existence of monomers and dimers in the mesoscopic protein-rich clusters was a central prediction of the kinetic model of cluster formation, put forth by Pan, *et al.* (32). Importantly, the demonstrated equilibrium between the clusters and the solution (33 38-40) enforces equal chemical potential of the monomer in the two phases and upholds the supersaturation that drives crystal nucleation in the clusters.

The role of dimers, likely bound by hydrophobic contacts, in the formation of the crystal nucleation precursors suggests that crystal nucleation can be controlled by transforming the interdomain interface to enhance or suppress its hydrophobicity. The volume of the crystal nucleation precursors can be enhanced, for instance, by removing the polar aminoacid residues, present at the interface between the structural domains of proteins incalcitrant to crystallization; such mutations may stabilize hydrophobically-bound domain-swapped dimers. This pathway to enhance nucleation is distinct from surface entropy reduction, in which charged aminoacid residues are removed from the protein surface (83, 84). Alternatively, chaotropic and cosmotropic agents that regulate hydrophobicity by modulating the water structures at non-polar protein surface patches may be applied to tune the dimer stability, and in this way control the volume of the cluster population and the rate of crystal nucleation (39, 40).

## AUTHOR CONTRIBUTIONS

V.L., P.G.V., and S.J.B. conceived this investigation. M.C.B., M.S.S., J.W.M., L.A.A., and D.H.H. performed experiments. L.A.A., S.J.B., V.L., J.C.C., and P.G.V. designed experiments. M.C.B. and V.L. analyzed the data. M.C.B. and P.G.V. wrote the manuscript, with contributions from V.L. and J.C.C.

## ACKNOWLEDGEMENTS

We thank M. Vorontsova-Kaissaratos for graphics help and K. Kourentzi and R.C. Willson for providing the hen-egg white lysozyme antibody. We thank the community of developers behind software libraries for Inkscape 0.91, Python (version 2.7.12) and the SciPy (version 0.18.1). This work was supported by NASA (Grants NNX14AE79G and NNX14AD68G) and NSF (Grant MCB-1518204).

